# Crosslinker Architectures Impact Viscoelasticity in Dynamic Covalent Hydrogels

**DOI:** 10.1101/2024.05.07.593040

**Authors:** Yung-Hao Lin, Junzhe Lou, Yan Xia, Ovijit Chaudhuri

## Abstract

Dynamic covalent crosslinked (DCC) hydrogels represent a significant advance in biomaterials for regenerative medicine and mechanobiology. These gels typically offer viscoelasticity and self-healing properties that more closely mimic *in vivo* tissue mechanics than traditional, predominantly elastic, covalent crosslinked hydrogels. Despite their promise, the effects of varying crosslinker architecture – side chain versus telechelic crosslinks – on the viscoelastic properties of DCC hydrogels have not been thoroughly investigated. This study introduces hydrazone-based alginate hydrogels and examines how side-chain and telechelic crosslinker architectures impact hydrogel viscoelasticity and stiffness. In hydrogels with side-chain crosslinking (SCX), higher polymer concentrations enhance stiffness and decelerates stress relaxation, while an off-stoichiometric hydrazine-to-aldehyde ratio leads to reduced stiffness and shorter relaxation time. In hydrogels with telechelic crosslinking, maximal stiffness and slowest stress relaxation occurs at intermediate crosslinker concentrations for both linear and star crosslinkers, with higher crosslinker valency further increasing stiffness and relaxation time. Our result suggested different ranges of stiffness and stress relaxation are accessible with the different crosslinker architectures, with SCX hydrogels leading to slower stress relaxation relative to the other architectures, and hydrogels with star crosslinking (SX) providing increased stiffness and slower stress relaxation relative to hydrogels with linear crosslinking (LX). The mechanical properties of SX hydrogels are more robust to changes induced by competing chemical reactions compared to LX hydrogels. Our research underscores the pivotal role of crosslinker architecture in defining hydrogel stiffness and viscoelasticity, providing crucial insights for the design of DCC hydrogels with tailored mechanical properties for specific biomedical applications.

## 1. Introduction

Mechanotransduction, the dynamic process by which cells translate mechanical cues into biochemical signals, has become a focal point of research over the past decade. This burgeoning interest stems from significant advancements in biomaterial design, fluorescent microscopy, and molecular biology tools, propelling our understanding of the intricate molecular pathways involved. Historically confined to the simplistic environment of petri dishes, the study of cellular behaviors has evolved, adopting specially treated surfaces and 2D cultures atop biomaterial hydrogels, and progressing to 3D culture of cells within engineered hydrogels with tailored properties^[1]^. This evolution marks a critical step towards accurately mimicking the cellular microenvironment, mirroring both the physical constraints and the complex composition of the extracellular matrix (ECM) ^[2–7]^. The importance of these advancements is underscored by a growing body of literature and clinical evidence, which suggests that variations in ECM stiffness or viscoelasticity serve as indicators and drivers of disease progression^[8–14]^. It is understood that biological tissues are viscoelastic^[15,16]^, and that the viscoelastic properties of the matrix critically influence cellular behaviors, including cell spreading^[2,17,18]^, cell migration^[19,20]^, ECM secretion^[17,21,22]^, stem cell differentiation^[14,23–25]^, morphogenesis^[26,27]^, cell cycle progression^[28,29]^, and even immune cell phenotypic changes^[30]^. Consequently, the rational design of biomaterial hydrogels, aimed at precisely modulating stiffness and viscoelasticity, has emerged as a cornerstone in the fields of regenerative medicine and mechanobiology.

Among the myriad of biomaterials, dynamic covalent crosslinked (DCC) hydrogels, particularly those utilizing Schiff-base chemistries including hydrazone, imine, and oxime bonds for crosslinking, have garnered attention for their injectability^[31–35]^, self-healing^[36–39]^, and viscoelastic nature^[17,18,24,26,40–43]^. The hydrazone linkage^[44,45]^, a reversible bond formed between hydrazine and aldehyde, is renowned for its biocompatibility and rapid kinetics, establishing it as a frontrunner in DCC chemistry for both regenerative medicine applications and mechanobiology investigations^[46]^. Researchers have develop various strategies to modify either synthetic or natural polymers for hydrazone crosslinked hydrogels, for example, polyethylene glycol (PEG)^[21,43,44]^, poly(L-glutamic acid) (PLGA)^[47]^, alginate^[40,48,49]^, xanthan^[37]^, hyaluronic acid (HA)^[17,32]^, and elastin-like protein^[24,33,50,51]^. Numerous studies have investigated how different parameters modulate the mechanical properties of the hydrazone hydrogels, such as polymer weight percentage^[17,50,52]^, molecular weight of the polymers^[34]^, degree of substitution (DS) of chemical groups on the polymers^[26,34,50,52]^, the stoichiometry of hydrazine to aldehyde ^[34,50]^, the relative ratio between aldehyde and benzaldehyde^[17,21,24,26,33]^, the valency of the crosslinker^[52]^, or the concentration of small molecule catalyst and competitor ^[32,33,52,53]^. We have recently elucidated that, among these parameters, the viscoelasticity of hydrogels is primarily determined by two factors: the exchange rate of crosslinks and the number of effective crosslinks per polymer chain^[52]^. However, the specific impact of crosslinker architectures—the polymeric structure of the crosslinks—on the stiffness and viscoelasticity of hydrazone hydrogels has not been thoroughly explored.

To elucidate how the structure of crosslinker regulates hydrogel mechanical properties, we developed alginate and PEG-based hydrazone crosslinked hydrogels formed from side-chain crosslinking (SCX) or telechelic crosslinking using crosslinkers of linear or star architectures (Figure 1A). SCX hydrogel occurs when alginate functionalized with hydrazine on the side-chain (AG-HYD) crosslinks with another alginate functionalized with aldehyde on the side-chain (AG-ALD). Telechelic crosslinking occurs when AG-HYD is crosslinked with aldehyde functionalized at the end-points PEG, including linear PEG-dialdehyde (PEG-2ALD) and “star” shaped 4-arm PEG-aldehyde (PEG-4ALD). The lack of cell adhesion binding site on either alginate or PEG offers an unbiased approach to study pure mechanistic effects on cellular behavior without the presence of biochemical cues. Most *in vitro* 3D cell culture systems using biopolymer hydrogels or reconstituted ECMs have a stiffness range from 100 Pa to 100 kPa^[2,4,5,9,15,21,24,27]^. Extensive studies have been performed using SCX hydrazone hydrogels. However, most of these gels do not exhibit physiologically relevant viscoelasticity (i.e., stress relaxation half-times of **τ**_1/2_ = 10-1000 sec) (Table. 1), unless an interpenetrating networks (IPNs) of collagen is incorporated to provide faster mode of relaxation^[17,18]^. The telechelic crosslinker architectures have gained popularity due to the ease of using small molecule adipic acid dihydrazide as a crosslinker^[40,42,47,49]^. However, few studies report the range of stiffness and viscoelasticity accessible using hydrogels with linear crosslinking (LX), and none indicate the range relevant for hydrogels with star crosslinking (SX)^[37]^. Another architecture star-star crosslinking, where hydrazine-modified multi-arms PEG crosslinked with aldehyde-modified multi-arms PEG, was not investigated in our work, but data from previous literatures were provided as comparison. Physiologically relevant stress relaxation timescale could only be achieved with either IPNs or extremely high polymer concentrations^[43,44,54]^.

**Table 1.**
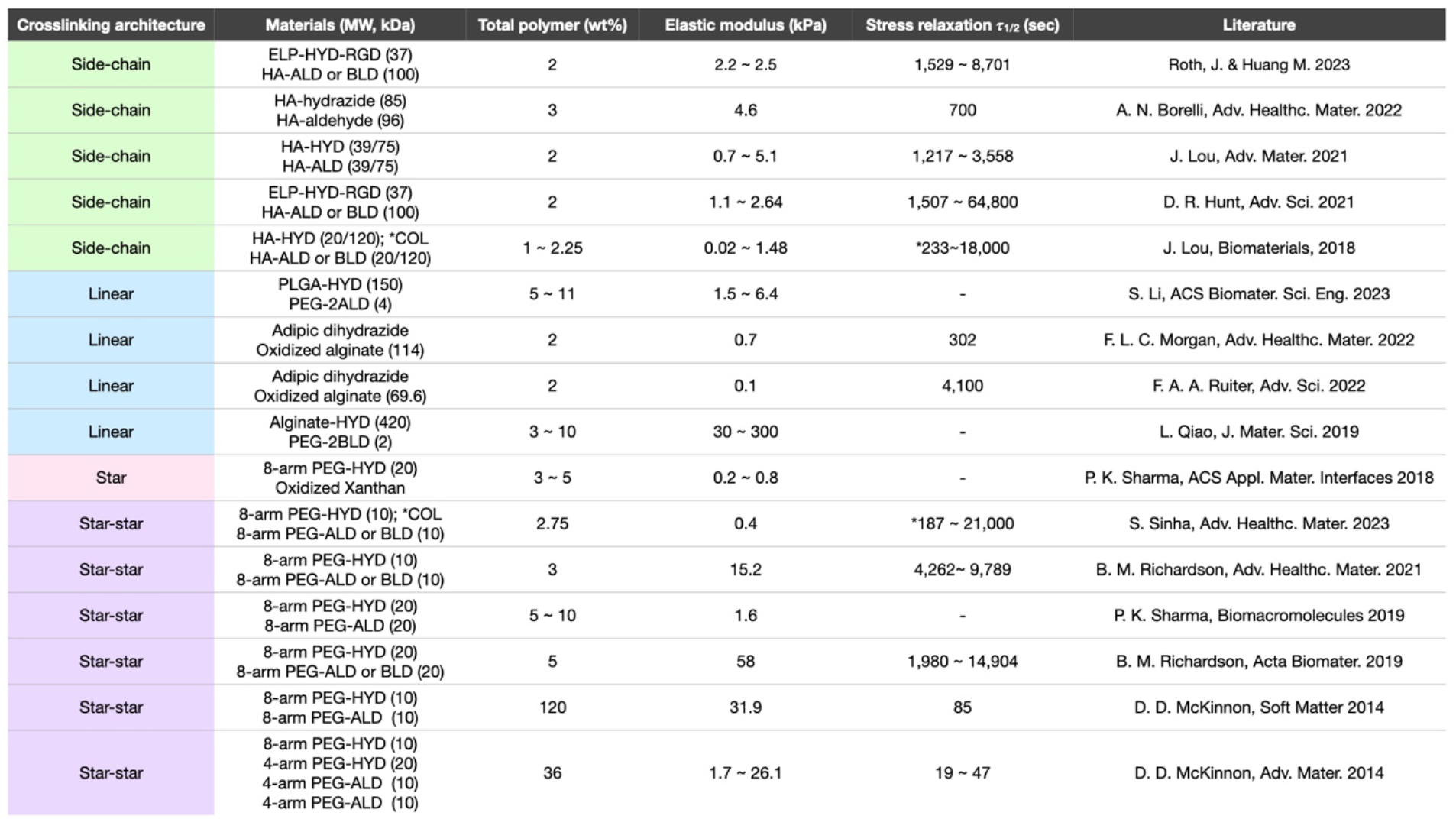
Stiffness and stress relaxation measured in previous studies using hydrazone-crosslinked hydrogels of various crosslinker architectures. ELP = elastin-like protein; HYD = hydrazine; RGD = arginyl-glycyl-aspartic acid; HA = hyaluronic acid; ALD = aldehyde; BLD = benzaldehyde; COL = collagen; PLGA = poly(lactic-*co*-glycolic acid); PEG-2ALD = polyethylene glycol dialdehyde. The asterisks indicate the fast relaxation were achieve with the incorporation of collagen as interpenetrating networks (IPNs) with the polymers.

**Figure 1.**
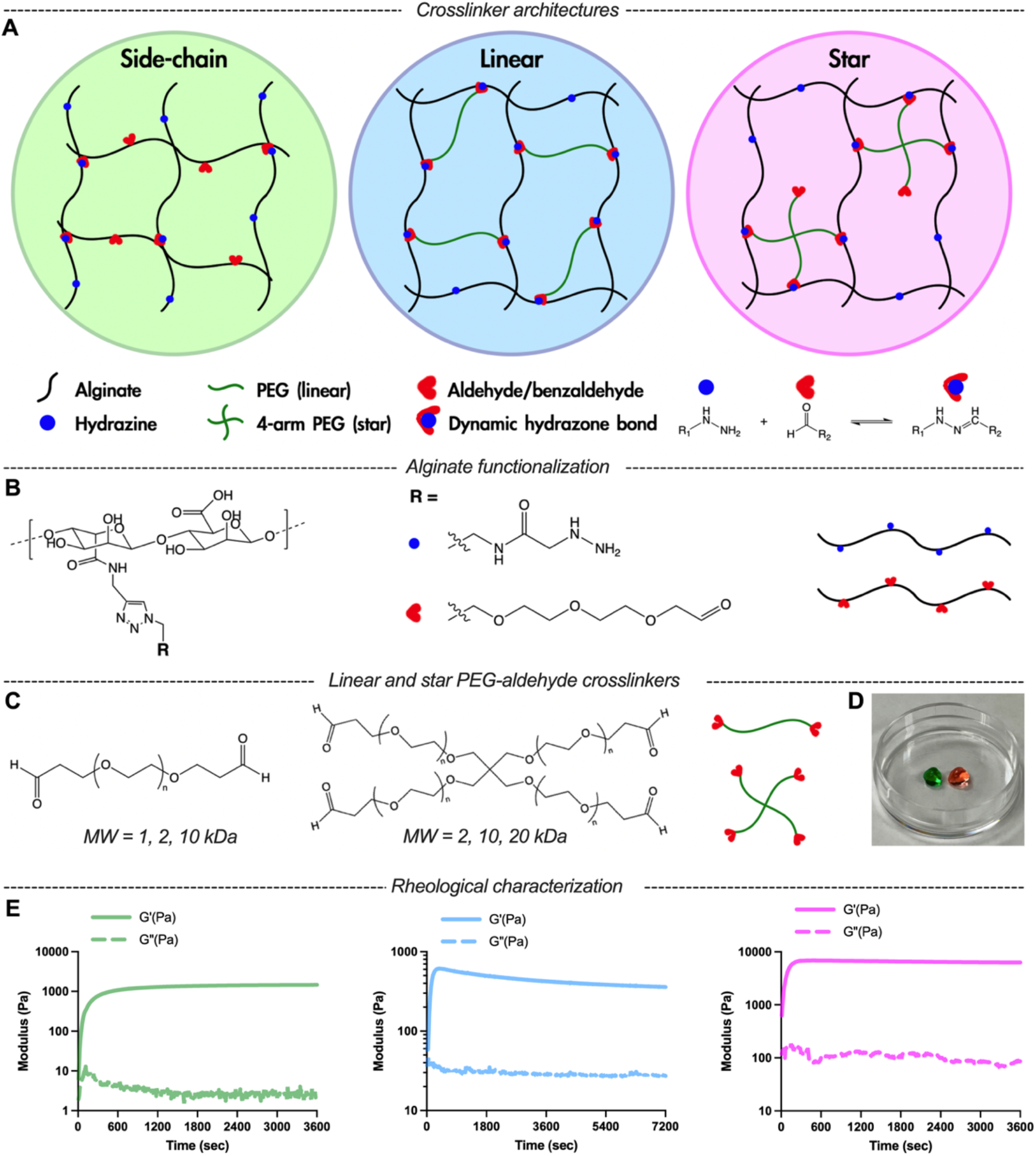
Dynamic covalent crosslinked (DCC) alginate hydrazone hydrogels can be formed with side-chain crosslinking (SCX), linear crosslinking (LX), and star crosslinking (SX) architectures. A) Schematics of SCX or telechelic crosslinking with LX or SX architectures for DCC hydrogels. B) Alginate was functionalized with either hydrazine or aldehyde groups. C) Linear and star PEG-aldehyde crosslinker molecules used for telechelic crosslinking. D) Representative image of two SX hydrogels placed in 6 mm cell culture dish, colored with two food dyes. E) Time evolution of storage and loss modulus of hydrogels with SCX (left), LX (center), and SX (right).

Here we demonstrate that DCC hydrazone-based hydrogels with SCX, LX, and SX exhibit distinct ranges of stiffness and viscoelasticity. Polymer concentration and the stoichiometry of hydrazine to aldehyde modulated stiffness and stress relaxation time of SCX hydrogels. Crosslinkers with different MW and valencies impacted the stiffness and viscoelasticity of LX and SX hydrogels. Further, we report that the mechanical properties of SX hydrogels were not significantly altered by encapsulation of high cell density, whereas cell density did impact the mechanical properties of SCX and LX hydrogels, further highlighting the importance of crosslinker architecture. This work aims to serve as a guidance of polymer and crosslinker architecture selection tailored to biomedical applications where stiffness and viscoelasticity are critical aspects.

## 2. Results and discussion

### 2.1 Controlled Functionalization of Alginate with Hydrazine or Aldehyde via Click Chemistry

To elucidate how crosslinker architecture affects the viscoelastic properties of DCC hydrogels, we engineered alginate-based hydrogels featuring diverse crosslinker architectures. Alginate was chosen as backbone polymers for its ubiquity in biomedical applications and amenability to chemical modifications. We modified 10% of carboxylate groups on alginate polymers with hydrazine or aldehyde following our previously reported two-step click functionalization (Figure 1B; Figure S1 and S2, Supporting Information)^[17]^. This synthetic method ensured equivalent functionalization levels of hydrazine and aldehyde for AG-HYD and AG-ALD, respectively. Notably, this click chemistry strategy for aldehyde functionalization circumvents the polymer degradation observed when using oxidation to form aldehydes on alginate^[55]^. DCC hydrogels were formed *in situ* via the hydrazone bond formation under physiological conditions when mixing hydrazine-containing polymer with aldehyde-containing polymers (Figure 1A). SCX hydrogels were formed by mixing AG-HYD with AG-ALD. For telechelic crosslinking, an extensive array of commercially available aldehyde-functionalized PEG variants, differing in valency and molecular weight (MW), were employed (Figure 1C). LX hydrogels were formed by mixing AG-HYD with PEG-2ALD, and SX hydrogels were formed by mixing AG-HYD with PEG-4ALD. Two of the SX hydrogels dyed with different colors were used to demonstrate self-healing capability of DCC hydrogels (Figure 1D; Figure S8, Supporting Information). To ensure a fair comparison of viscoelastic properties across different architectures, the degree of substitution (DS) of hydrazine on AG-HYD was fixed, given its substantial influence on the hydrogels’ elastic modulus^[34]^.

### 2.2 Hydrogels with Side-chain Crosslinking Exhibit Slow Stress Relaxation

To investigate the mechanical properties of SCX hydrogels, we utilized a model system consisting of AG-HYD and AG-ALD, both at a MW of 28 kDa and a DS of approximately 10%. The use of SCX architecture offered several distinct advantages: the simplicity of employing identical polymer backbones modified with different chemical groups; the availability of a broad spectrum of biopolymers, primarily polysaccharides, with side-chain functional groups amenable to chemical modifications; and flexibility of tuning the degree of substitution for a range of crosslinking densities. The viscoelastic properties of SCX hydrogels were adjusted by varying the total polymer concentration and the stoichiometry of the hydrazine and aldehyde functional groups. Evaluation of mechanical properties was conducted through a series of rheological tests: a time sweep, frequency sweep, and a stress relaxation test (Figure 1E, 2A and 2B). As expected, at a 1:1 stoichiometry of AG-HYD to AG-ALD, an increase in total polymer concentration resulted in a higher elastic modulus, attributed to the greater amount of polymer and higher density of crosslinking sites (Figure 2C)^[17,56]^. Hydrogels with 1 wt% alginate exhibited an elastic modulus of 0.99 kPa, which increased to 17.85 kPa at a concentration of 4 wt%.^[16,49]^ Hydrogels formulated with off-stoichiometric ratio of hydrazine to aldehyde (either 1:3 or 3:1) demonstrated reduced stiffness and accelerated stress relaxation at the same overall polymer concentration (Figure 2F), aligning with the biphasic influence of stoichiometry on stiffness observed in prior studies^[34]^. However, the similar biphasic relationship on stress relaxation time has not been previously reported. Overall, SCX hydrogels presented substantial range of stiffness tunability, from 0.24 to 18 kPa across the 1 to 4 wt% gel formulations of various hydrazine to aldehyde ratio. However, the range of viscoelasticity was constrained in these gels. The stress relaxation half-times (*τ*_1/2_), the time required for stress in hydrogel to relax to half of its initial value, consistently exceeded 1,500 seconds (Figure 2E, H) and loss tangents remained below 0.03 (Figure 2D, G), which lie outside of the range relevant to most soft tissues^[16]^.

**Figure 2.**
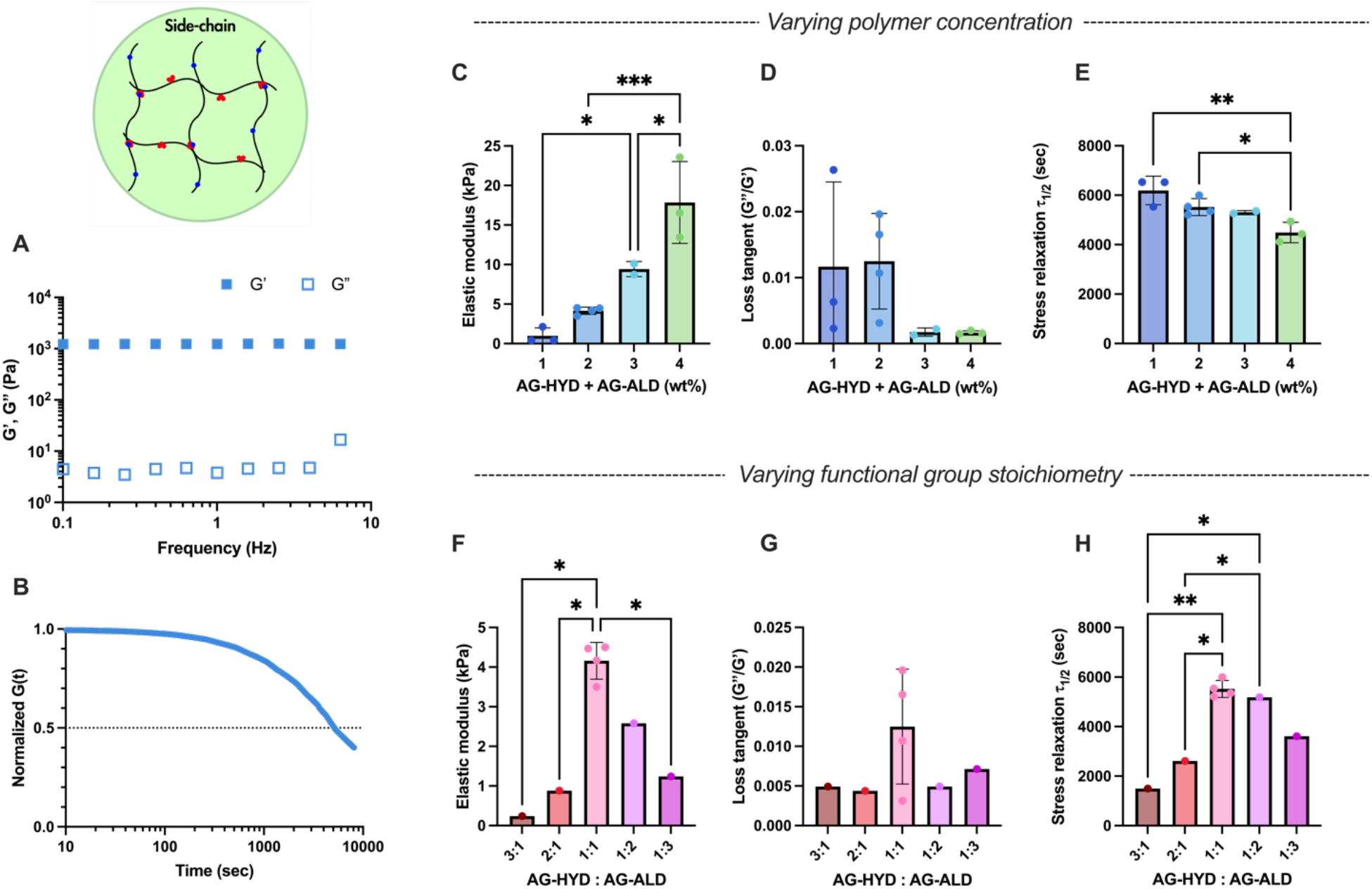
Rheological characterizations of SCX hydrogels. A-B) Representative of frequency sweep (average of n = 4) and stress relaxation (average of n = 3) tests using formulation of 2 wt% alginate at AG-HYD to AG-ALD ratio of 1:1. C-E) The elastic modulus, loss tangent, and stress relaxation half-times of hydrogels with varying alginate concentration under same AG-HYD to AG-ALD ratio of 1:1. F-H) The elastic modulus, loss tangent, and stress relaxation half-time of hydrogels with varying AG-HYD to AG-ALD ratios, all with the same alginate concentration of 2 wt%. One-way ANOVA with Tukey’s post-hoc test. * *P* ≤ 0.05, ** *P* ≤ 0.01, \*\*\**P* ≤ 0.001; any statistical relationship not indicated means not significant.

### 2.3 Intermediate crosslinker concentration leads to maximum stiffness and stress relaxation time in hydrogels with telechelic crosslinking

We next tested the range of properties accessible for hydrogels with telechelic crosslinking, Including LX and SX. Similar to SCX hydrogels, we hypothesized that an intermediate ratio of hydrazine to aldehyde within telechelic crosslinked structures would yield the peak in both stiffness and stress relaxation half-time. Indeed, we found that in either LX (Figure 3A, B) or SX hydrogels (Figure 3C, D), the intermediate PEG-aldehyde crosslinker concentration leads to maximal stiffness and stress relaxation time under the same AG-HYD concentration. At low crosslinker concentrations, there are likely not sufficient levels of interchain crosslinking to form robust percolating polymer networks. Conversely, excessive aldehyde crosslinker would be expected to saturate the hydrazine groups on alginate chains, resulting in dangling PEG chains that do not contribute to the crosslinked network^[57]^. This higher polymer concentration does not contribute to higher stiffness, indicating the hydrazine to aldehyde ratio dominates the mechanical properties of the hydrogel instead of entanglement (Figure S7, Supporting Information). Overall, within telechelic crosslinker architectures, the intermediate concentration of crosslinker leads to peak stiffness with slowest stress relaxation.

**Figure 3.**
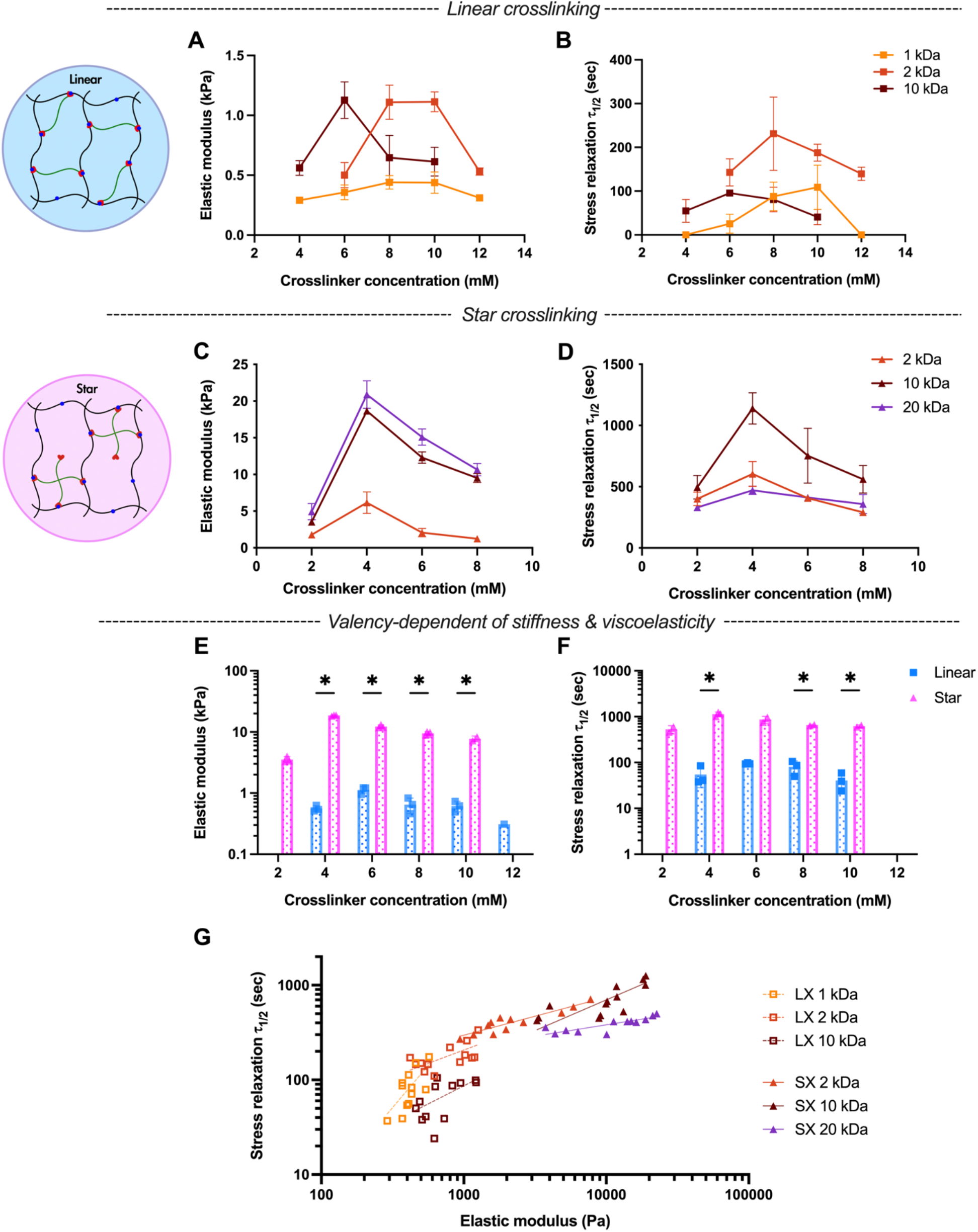
Rheological characterizations of hydrogels with telechelic crosslinking, using crosslinkers with either linear or star architectures. A-B) The elastic modulus and stress relaxation half-time of LX hydrogels and varying concentrations of PEG-dialdehyde crosslinkers of different molecular weight. C-D) The elastic modulus and stress relaxation half-time of SX hydrogels and varying concentrations of 4-arm PEG-dialdehyde crosslinker of different molecular weight. E-F) Elastic modulus and stress relaxation half-time of LX versus SX hydrogels. Multiple unpaired Welch’s t-test with Holm-Šídák multiple comparison. G) DCC alginate hydrazone hydrogels with telechelic crosslinking present correlation of stress relaxation half-time and elastic modulus. Linear regression lines were overlayed, dashed lines for LX hydrogels and solid lines for SX hydrogels. Pearson *r* coefficient and *P* value: linear 1 kDa (0.69, *), 2 kDa (0.62, *), and 10 kDa (0.59, *); star 2 kDa (0.92, ****), 10 kDa (0.76, **), and 20 kDa (0.82, **). * *P* ≤ 0.05, ** *P* ≤ 0.01, \*\*\**P* ≤ 0.001, \*\*\*\**P* ≤ 0.0001.

### 2.4 Modulation of Viscoelastic Properties by Crosslinker Molecular Weight and Valency in Telechelic Crosslinking

To further explore the parameters that could modulate the viscoelasticity of hydrogel with telechelic crosslinking, we delved into the roles of crosslinker MW and valency. We hypothesized that with telechelic crosslinking, under the same crosslinker molar concentration (mM), crosslinkers with higher MW may lead to higher stiffness and stress relaxation time due to potential entanglement associated with higher polymer weight concentrations. Surprisingly, crosslinkers of intermediate MW exhibited the higher levels of stiffness and stress relaxation, observable in both LX (Figure 3A, B) and SX (Figure 3C, D) hydrogels. This phenomenon may arise from the intermediate crosslinkers’ length aligning more closely with the spacing between available hydrazine groups in the networks, optimizing interchain crosslinking. Shorter crosslinkers likely fail to bridge two alginate chains, while longer crosslinkers may inadvertently facilitate intrachain crosslinking, subsequently reducing the overall hydrogel elasticity. Further, we hypothesized that, under the same crosslinker molar concentration and MWs, crosslinkers with higher valency (e.g., 4-arm for star) would yield more elastic hydrogels compared to those with lower valency (e.g. 2-arm for linear). Consistent with this expectation, SX hydrogels resulted in up to a 33-fold increase in stiffness and 20-fold increase in stress relaxation time relative to LX hydrogels (Figure 3E, F; Figure S9, Supporting Information). This agrees with the previous study that increasing the number of crosslinking sites per chain slows stress relaxation^[52]^. Notably, stress relaxation time exhibited a positive logarithmic correlation with elastic modulus across different crosslinker architectures (Figure 3G). Though, the slopes of these relationships did depend upon the crosslinker MW and valency. These findings underscore the pivotal influence of crosslinker MW and valency on the viscoelastic characteristics of DCC hydrogels.^18^

### 2.5 DCC hydrogel viscoelasticity is crosslinker architecture dependent

To elucidate whether crosslinker architectures could contribute to different range of viscoelasticity of DCC hydrogel, we overlayed all the measurements collected in this work on stress relaxation versus stiffness plot with colors indicating the loss tangent (Figure 4A). Strikingly, we observed distinct regions occupied by hydrogels with side-chain, linear, and star crosslinking, with minimal overlap. LX hydrogels lie in a regime of low stiffness, fast stress relaxation, and high loss tangent. SCX hydrogels reside in a region characterized by slow stress relaxation and minimal loss tangent. SX hydrogels exhibit higher stiffness but have a lower loss tangent and slower stress relaxation compared to LX. The distinct regions occupied by hydrogels with different crosslinker architectures became more apparent when we plotted stress relaxation half-time against loss tangent (Figure 4B). Data from existing literature were consistent with our results, though less data existed for LX and no data was found for SX (Figure 4C). Overall, SCX hydrogels exhibit a broad stiffness range but predominantly fall in a slow relaxation regime (>4,000 sec). Achieving faster stress relaxation requires a high off-stoichiometric ratio of hydrazine to aldehyde, which significantly reduces stiffness (Figure 2F, H). In contrast, LX hydrogels demonstrate much faster stress relaxation, akin to biological tissues, but with a limited stiffness range of 0.3 to 1.1 kPa (Figure 3A, B)^[16]^. SX hydrogels, however, offer a wide stiffness spectrum while maintaining fast stress relaxation close to the physiological relevant range. Overall, these findings underscore the unique viscoelastic ranges presented by DCC hydrazone hydrogels of varied crosslinker architectures. Further and importantly, understanding the interplay between various crosslinking parameters can guide the design of studies designed at elucidating how one mechanical property of the hydrogels independently regulates cellular behaviors of interest. For instance, to investigate the mode of cell migration in DCC hydrogels of same stiffness but different stress relaxation time^[19]^, hydrogels that follow a vertical line in Figure 4A can be utilized.

**Figure 4.**
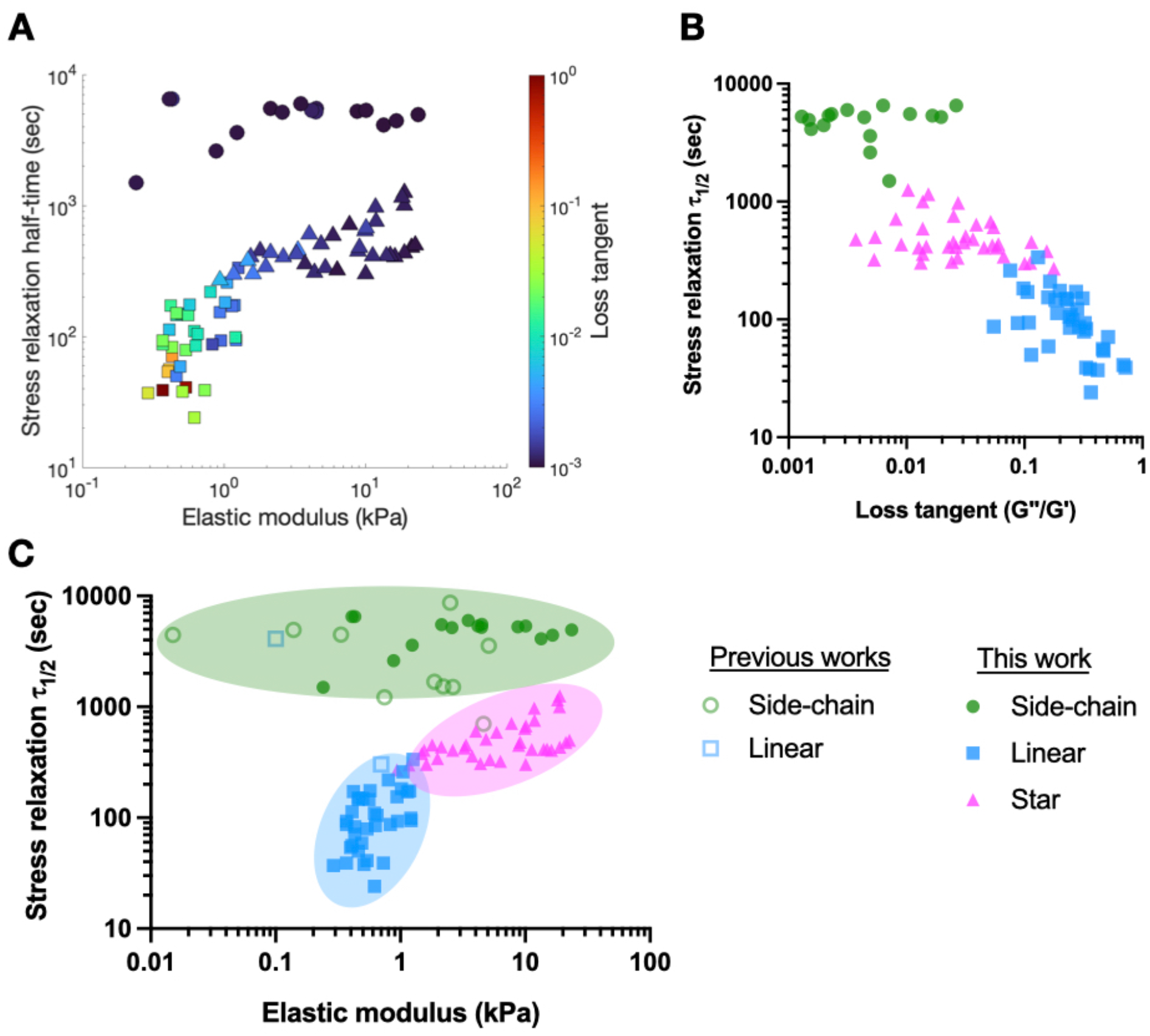
DCC hydrogel stiffness and viscoelasticity are crosslinker architecture dependent. A) The elastic modulus, stress relaxation half-time, and loss tangent of hydrazone hydrogels with SCX, LX, and SX collected in this work. B) The stress relaxation half-time and loss tangent of hydrazone hydrogels with SCX, LX, and SX collected in this work. C) Combined data from previous works and this work demonstrate hydrazone hydrogels with SCX, LX, and SX present distinct range of stiffness and stress relaxation half-time with minimal overlaps. SCX (circle), LX (square), and SX (triangle).

### 2.6 Impact on Mechanical Properties from Encapsulated Cells Depends on Crosslinker architectures

The impact of encapsulated cells on the viscoelastic properties of hydrogels was assessed to gauge the use of these materials for 3D cell culture or cell and drug delivery applications. Given the marked increase in stiffness and stress relaxation time from SX compared to LX hydrogels, we speculated that the additional valency of SX might enhance resistance to competitive chemical reactions that threaten the integrity of hydrazone bonds. Such competitive interactions may arise from glucose in its open-chain form found in cell culture media, as well as free amines from amino acids, serum, cell membrane proteins, and secreted ECM components^[58,59]^. We focused on hydrogels with the highest identified stiffness and stress relaxation time for both linear and star crosslinking architectures, as shown in Figure 3A and Figure 3C: 2wt% AG-HYD with 8 mM PEG-2ALD (2 kDa) for LX hydrogels and 2wt% AG-HYD with 4 mM PEG-4ALD (10 kDa) for SX hydrogels. In LX hydrogels, encapsulating 10 million MDA-MB-231 breast cancer cells per milliliter resulted in an approximate 70% decrease in initial stiffness and a 40% reduction in stress relaxation time (Figure 5A, B). Conversely, SX hydrogels maintained their stiffness and exhibited only a minor reduction in stress relaxation time under identical conditions of AG-HYD concentration and cell density, showcasing the valency-dependent resistance of hydrazone bonds against competitive reactions. Interestingly, SCX hydrogels at the same alginate concentration also demonstrated a 30% reduction in stiffness. However, SCX hydrogels with 6wt% alginate, possessing a polymer content comparable to the formulation of SX hydrogels, did not show compromised stiffness, though its stress relaxation time was halved upon cell encapsulation (Figure S10, Supporting Information). This suggests that with a given crosslinker architecture, the stability of hydrazone bonds against competitive reactions may also be influenced by the crosslink density associated with the increased polymer content. These observations underscore the critical need for careful selection of crosslinker architecture, especially when alterations in mechanical properties due to encapsulated entities are of concern in applications. For example, in hydrogel-based cell, gene, and drug delivery systems, the influence of payload density on mechanical properties and release kinetics may require careful study.

**Figure 5.**
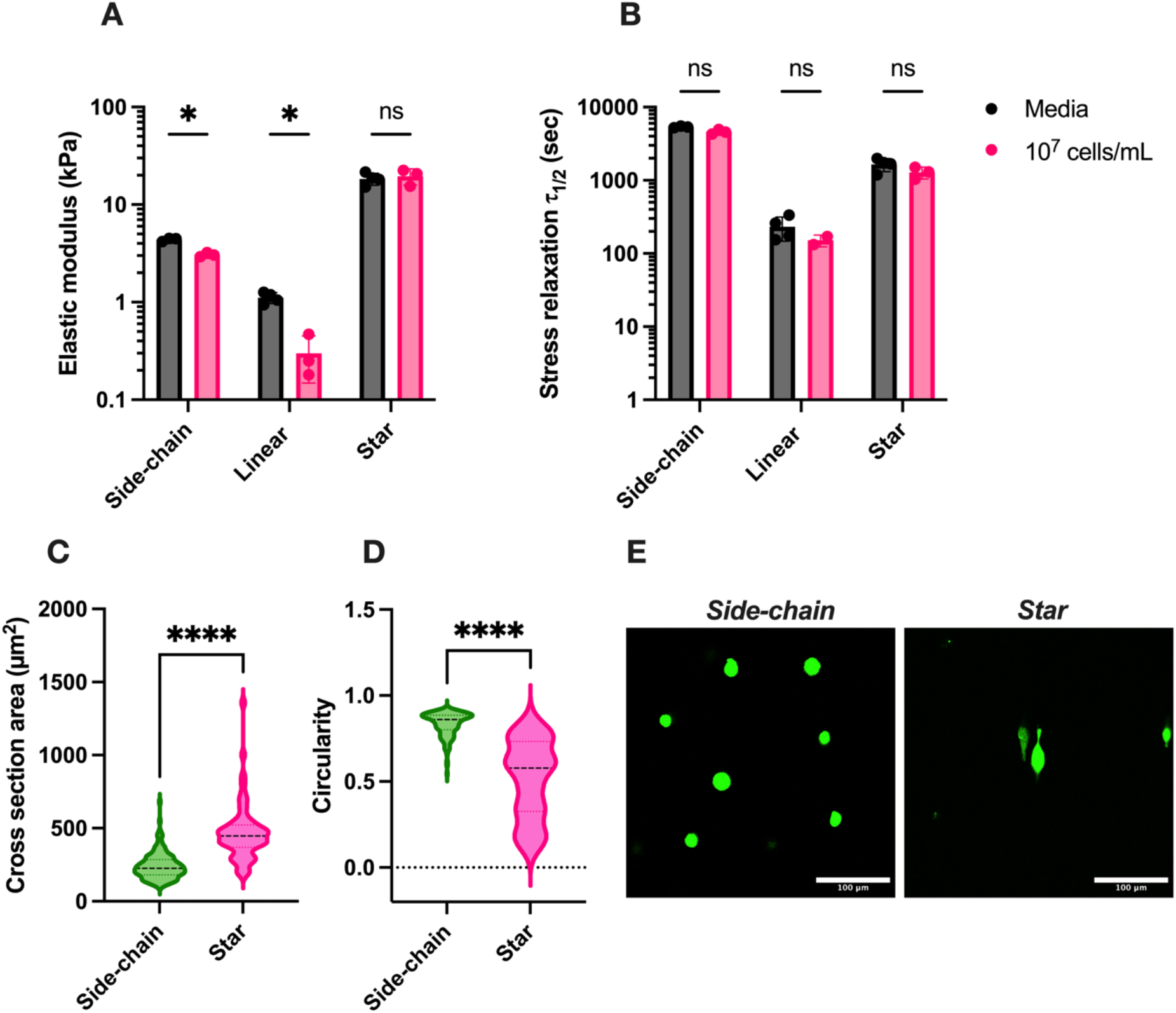
3D cell culture and impact of cell encapsulation on hydrogel mechanical properties. A-B) The impact of encapsulating 10 million cells per milliliter in side-chain, linear, and star crosslinking gels on stiffness and stress relaxation half-time. Multiple unpaired Welch’s t-test with Holm-Šídák multiple comparison. C-D) The cross-section area and circularity of MDA-MB-231 cells encapsulated in SCX and SX hydrogels of similar stiffness (17.8 vs 18.7 kPa) but varied stress relaxation time (4,492 vs 1,139 sec). n=148 for SCX and n=62 for SX. Two-tailed unpaired t-test. E) representative images of cell morphology in SCX and SX hydrogels, stained with Calcein-AM dye in green. Scale bar = 100 μm. ns, not significant; * *P* ≤ 0.05, \*\*\*\**P* ≤ 0.0001.

### 2.7 3D cell culture in hydrogels of varied crosslinker architecture

Next, we explored 3D cell culture in hydrogels with different crosslinker architectures. After 24 hours of culture, the cross-sectional area and circularity of MDA-MB-231 cells were analyzed. In SCX hydrogels (E=17.8 kPa, *τ*_1/2_=4,492 sec), cells predominantly adopted a rounded morphology, characterized by a mean circularity of 0.835 and a cross-sectional area of 241.3 μm^2^ (Figure 5C, D). In contrast, cells in SX hydrogels (E=18.7 kPa, *τ*_1/2_=1,139 sec) displayed an ellipsoidal shape, with a decreased mean circularity of 0.513 and an increased cross-sectional area of 468.2 μm^2^. This shows that under the same stiffness, the hydrogel with faster stress relaxation leads to more elongated cell morphology (Figure 5E). LX hydrogels, however, experienced rapid disintegration within 24 hours in cell culture media, precluding any cell shape and area analysis. This fast erosion is likely due to the reversible nature of hydrazone bonds, the limited valency of PEG-dialdehyde crosslinkers, and competitive interactions from free amines targeting imine bond formation. To improve the stability of linear crosslinked hydrogels in cell culture conditions, strategies such as increasing the DS of AG-HYD or incorporating PEG-dibenzaldehyde for more durable hydrazone linkages may be necessary^[17,21]^.

## 3. Conclusion

In this work, we demonstrate the influences of DCC hydrazone crosslinker architectures –– whether side-chain or telechelic––on viscoelasticity and stiffness of the resulting hydrogel. We have identified that SCX yields hydrogels with a considerable range of stiffness but consistently slower stress relaxation. In contrast, telechelic crosslinking, particularly with SX, affords a versatile modulation of hydrogel mechanical properties, enabling the achievement of higher stiffness than LX counterparts and more rapid stress relaxation in comparison to SCX hydrogels. Furthermore, we elucidate that SX hydrogels exhibit remarkable mechanical stability with high encapsulating cell density. These findings underscore the importance of crosslinker architecture consideration in the design of biomaterial hydrogels tailored for specific biomedical applications.

## 4. Experimental Section

### Alginate modification

#### Alginate-alkyne (AG-ALK)

The alginate modification scheme was adapted from a previous study that reported the coupling of alkyne and hydrazine groups to hyaluronic acid^[17]^ (Fig. S1). Low molecular weight alginate (#4200501, batch BP-1212-24, PRONOVA, UP VLVG) (1 g, 5.7 mmol, 1 eq.) was fully dissolved in 100 mL MES buffer (200 mM MES hydrate, 154 mM NaCl, pH=4.5). The following chemicals were then added quickly to the 1 wt% alginate solution with the following order: N-hydroxysuccinimide (NHS) (261.6 mg, 2.3 mmol, 0.4 eq., #130672 Sigma Aldrich), EDC (435.5 mg, 2.27 mmol, 0.4 eq., #E6383, Sigma Aldrich), then propargylamine (145.6 uL, 2.273 mmol, 0.4 eq., #P50900, Sigma Aldrich). After adjusting pH of the mixture to 6, the solution was stirred at room temperature for 4 hrs, followed by quenching with hydroxylamine hydrochloride (631.7 mg, 9.09 mmol, 4 eq., #159417 Sigma-Aldrich) for 30 min. The solution was dialyzed (MWCO = 10 kDa) in deionized (DI) water at 4°C, with two water exchange in the first hour and 8 more times in the following 3 days. The solution was lyophilized (Labconco freezone 4.5 lyophilizer) for 5 days to get white cotton-like alginate-alkyne (AG-ALK) product. The AG-ALK product was stored at −30°C until use.

#### Alginate-hydrazine (AG-HYD) and Alginate-aldehyde (AG-ALD)

AG-ALK (500.0 mg, 2.347 mmol, 1 eq.) was dissolved in 25 mL of 0.1M PBS (10 mM NaH2PO4, 9 mM Na2HPO4, 0.1 M NaCl, pH=7.4), then N-(3-azidoethyl)-2-hydrazineylacetamide (compound **2**, 296.71 mg, 1.878 mmol, 0.8 eq.) or azido-aldehyde (compound **3**, 424.41 mg, 1.878 mmol, 0.8 eq.) was added to the alginate solution. Sodium ascorbate (1045.8 mg in 1.5 mL DI water, #11140 Sigma-Aldrich), copper sulfate pentahydrate (176.1 mg in 1.5 mL DI water, #209198 Sigma-Aldrich), and alginate solution were sealed in separated vials and bubbled with nitrogen for at least 30 min. Sodium ascorbate (200 uL; 139.4 mg, 0.704 mmol, 0.3 eq.) and copper sulfate pentahydrate solutions (100 uL; 11.74 mg, 0.047 mmol, 0.02 eq.) were added to the alginate solution sequentially under continuing nitrogen bubbling. The alginate solution was stirred for 1 day at room temperature (or 2 days at 4°C for AG-ALD) while protected from light. The reaction was quenched with 0.5 wt% EDTA (10 mL, 0.4x v/v to alginate solution) for 30 min at room temperature. The solution was dialyzed (MWCO = 10 kDa) in DI water at 4°C with 10 buffer exchanges in 3 days, then filtered through 0.22 μm filter. The filtered solution was lyophilized for 5 days to get white cotton-like AG-HYD or AG-ALD product. The final lyophilized product was stored at −30°C until use. The degree of substitution (DS) on alginate was determined using NMR spectroscopy under 70°C by integration of the proton signal on triazole relative to the proton on the first carbon of guluronic acid on alginate backbone and assumed guluronic acid to mannuronic acid ratio of 1.5. 1H NMR integration indicated that ∽10% of the carboxylate groups on the alginate backbone have been functionalized with either hydrazine or aldehyde. Triazole H: ^1^H NMR (500 MHz, D_2_O) *δ* 7.88 (s, 1H). Guluronic acid H: ^1^H NMR (500 MHz, D_2_O) *δ* 5.03 (s, 1H).

### Hydrogel preparation

AG-HYD, AG-ALD, and PEG-aldehyde of various molecular weight and valencies were dissolved in low glucose Dulbecco’s modified Eagle’s medium (DMEM-LG) (#11885084, Gibco) with 25 mM of HEPES and adjust pH to 7.4∽7.5, monitored by micro pH probe (pH Sensor InLab® Micro, #51343160, METTLER TOLEDO). It has been shown that free amines, either from amino acids or proteins, may compete with the hydrazone bonds via imine bond formation with free aldehydes ^[58,59]^. Thus, we intentionally excluded fatal bovine serum and antibiotics from the low glucose Dulbecco’s Modified Eagle Medium (DMEM-LG) to reconstitute the AG-HYD and AG-ALD polymers when performing rheological tests. Due to the pH sensitivity of hydrazone bond formation, we supplemented 25 mM of HEPES in the media to help with buffering the pH when CO_2_ control was not feasible under the shear rheometer. The pH of the reconstituted polymer was monitored and adjusted to neutral using micro pH meter before each measurement to minimize the artifact of changing stiffness and stress relaxation time results from the gradually rising pH in atmospheric condition. By mixing AG-HYD with either AG-ALD or PEG-aldehyde in eppendorf, hydrogels were rapidly formed after few seconds of vortexing. For example, to prepare a 100 uL of 2wt% SCX hydrogel (HYD: ALD = 1 : 1), 50 uL of 4wt% AG-ALD was first mixed with 25 uL of media. Then 50 uL of 4wt% AG-HYD was added to the mixture. To prepare a 100 uL of 2wt% SX hydrogel with 4 mM of PEG-4ALD, 16 uL of 25 mM PEG-4ALD was first mixed with 34 uL of media. Then 50 uL of 4wt% AG-HYD was added to the mixture.

### Rheological characterization

Rheology measurements were performed on Discovery HR-2 hybrid rheometer (TA Instruments) to quantify the frequency-dependent viscoelastic properties including storage modulus, loss modulus. The rheometer was heated to 37°C before measurements. 60 uL of hydrogels were deposited directly onto the rheometer base plate. An 8-mm flat plate was then immediately lowered to contact the gel to form an 8-mm disk. Mineral oil (Sigma) was applied to seal the edges of the gel to prevent hydration. Time sweep was performed at 1 rad s^-1^ and 1% strain until storage and loss modulus plateaued. Assuming Poisson’s ratio (ν) of 0.45, the elastic modulus was calculated based on the equation:

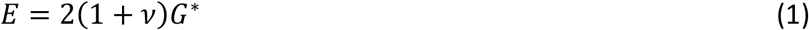

were the complex modulus (G^*^) was derived from the measured storage (G’) and loss modulus (G”) by:

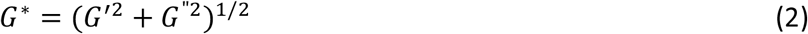

Frequency sweep measurements were then performed from 0.1 to 100 rad s^-1^ at 1% strain. For stress relaxation measurements, after letting hydrogel sit for 5 min to equilibrate, a constant strain of 15% was applied to the gel and the resulting stress was recorded. The stress relaxation time (*τ*_1/2_) was defined as the time that resulting stress decayed to one-half of the initial value. The mechanical characterization from previous works were analyzed by an online data-capturing program WebPlotDigitizer (Rohatgi, WebPlotDigitizer, 2022).

### Encapsulation of cells in hydrogel

MDA-MB-231 cells were cultured in high glucose Dulbecco’s modified Eagle’s medium (#11995065, Gibco) containing 10% fetal bovine serum (#35-015-CV, Corning) and 1% penicillin/streptomycin (#15140122, Gibco). Cells were passaged around 70% confluency, spun down, resuspended in DMEM-LG containing 10% fetal bovine serum and 1% penicillin/streptomycin. Cell density was measured using Vi-CELL cell viability analyzer (Beckman Coulter). To make a cell-containing hydrogel for mechanical characterization, cell suspension was added to the mixture of media and AG-ALD or PEG-aldehyde, before mixing with AG-HYD. The final cell density in the hydrogels were 10^7^ cells/mL. For 24 hours of 3D cell culture, MDA-MB-231 cells were encapsulated at 10^6^ cells/mL. 100 μL of cell-containing hydrogels were added to the bottom of a well in 8-well LabTek chamber (#155360, Thermo Scientific). After 30 minutes, 100 uL of cell culture media was added on top of each gel, then the hydrogels were incubated in incubator for 24 hours. Calcein-AM dye (#C1430) of 1:1000 dilution was then used to stain cells for 30 minutes in the incubator before imaging. The formulation of side-chain crosslinking hydrogel was 2wt% of alginate with AG-HYD to AG-HYD ratio of 1:1. The formulation LX hydrogel was 2wt% AG-HYD with 8 mM of 2 kDa PEG-dialdehyde. The formulation of SX hydrogel was 2wt% of AG-HYD with 4 mM of 10 kDa 4-arm PEG-aldehyde.

### Confocal microscopy

Microscopy was performed on a laser-scanning Leica SP8 confocal microscope with 25x 0.95-NA water immersion objective. Cell morphologies in 3D culture were recorded by confocal imaging and the cross-section area and circularity of cells were analyzed on Fiji ImageJ.

### Statistical analysis

Statistical analyses were performed using GraphPad Prism 10 software. Data are presented as mean ± SD if not otherwise mentioned. One-way analysis of variance with Tukey’s post-hoc test was used to analyze the elastic modulus, loss tangent, and stress relaxation half-time of SCX hydrogels (Fig. 2C-H), and the effects of entanglement (Figure S7, Supporting Information). Unpaired Welch’s t-test with Holm-Šídák post-hoc test was used to compare the mechanical properties of LX and SX hydrogels (Fig. 3E, F and Figure S9, Supporting Information), and cell encapsulation effects on mechanical properties comparing three different crosslinker architectures (Fig. 5A, B). Two-tailed Pearson correlation was used for the correlations between stress relaxation half-time and elastic modulus of hydrogels with telechelic crosslinking (Fig 3G). Two-tailed unpaired t-test was used to compare cell circularity and cross-sectional area of SCX and SX hydrogels (Fig. 5C, D) and cell encapsulation effects on mechanical properties of SCX hydrogels at 6wt% alginate (Figure S10, Supporting Information). In all the cases, P ≤ 0.05 was considered to be statistically significant. All significance levels are denoted as follows: *P ≤ 0.05, **P ≤ 0.01, ***P ≤ 0.001, ****P ≤ 0.0001 and ns indicates not significant.

## Supporting information

Supporting Informations

## 5. Author contributions

Y.H.L., Y.X., and O.C. conceived the ideas and designed experiments; Y.H.L conducted experiments and analyzed data; Y.H.L. and J.L. designed the synthesis schemes; Y.X. and O.C. provided supervision and oversight to the work; Y.H.L and O.C. wrote and edited the manuscript; J.L. and Y.X. reviewed the manuscript.

## 6. Acknowledgements

O.C. gratefully acknowledges funding support for this work from the NIH grants R01 AR081993 and R37 CA214136.

## 8. Data Availability Statement

The data in this study are available from the corresponding author upon reasonable request.

