## Supporting Informations for "Crosslinker Architectures Impact Viscoelasticity in Dynamic Covalent Hydrogels"

**Characterizations.**

Proton NMR ($^1$H NMR) spectra were obtained with Inova 300 and Inova 500 spectrometers. The chemical shifts are presented in parts per million (ppm) and were calibrated using the minor proton-containing component of the solvent as the internal reference. Specifically, for $CDCl_3$, the reference chemical shift was at 7.26 ppm, while for $D_2O$, the shifts were recorded at 4.77 ppm at 20°C and 4.28 ppm at 70°C.

**Synthesis of small molecules.**

*Synthesis of compound **1**: 2-azidoethylamine*

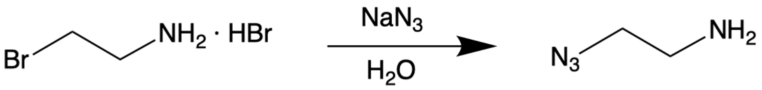


2-Bromoethylamine hydrobromide (10.1 g, 49.29 mmol, 1 eq.) and sodium azide (8.01 g, 123.24 mmol, 2.5 eq.) were dissolved in 30.3 mL DI water. The mixture was stirred at 90°C in oil bath for more than 18 hrs while protected from light. After the mixture was cooled down to room temperature, pH was adjusted to greater than 11 using 10N NaOH, then was stirred for 30 minutes. Extract the product with ethyl ether for three times then dry with sodium sulfate. The excess solvent was removed by rotavap, and the residue was final product as a brownish oil (1.46 g, 34% yield). $^1$H NMR (300 MHz, $CDCl_3$) δ 3.37 (t, $J$ = 5.7 Hz, 2H), 2.88 (t, $J$ = 5.7 Hz, 2H)

*Synthesis of compound **2**: N-(3-azidoethyl)-2-hydrazineylacetamide*

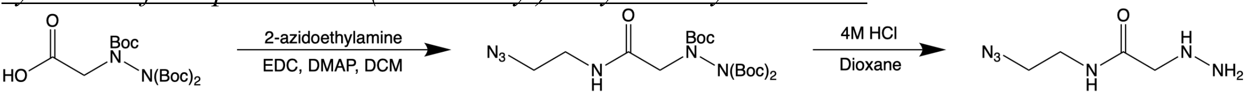


Tri-Boc-hydrazinoacetic acid (3.5 g, 8.964 mmol, 1 eq.), 2-azidoethylamine (compound **1**, 0.848 g, 9.861 mmol, 1.1 eq., #68972 Sigma Aldrich), 1-ethyl-3-(3-dimethylaminopropyl) carbodiimide (EDC) (2.233 g, 11.654 mmol, 1.3 eq.), and 4-dimethylaminopyridine (DMAP) (0.219 g, 1.793 mmol, 0.2 eq.) were dissolved in 12 mL dichloromethane (DCM). The solution was stirred at room temperature for 1 day, then diluted with DCM, washed with brine twice, and dried with sodium sulfate. The solvent was removed under reduced pressure, and the residue was purified by silica gel chromatography (ethyl acetate/hexane, 3:1 v/v, to 2:1, to 1:1) to obtain the Boc-protected product as a colorless oil (4.03 g, 98% yield). The isolated Boc-protected product was then dissolved in 10 mL 4 M HCl in dioxane. After stirring at room temperature for 4 h, the product precipitated as a sticky yellow solid was centrifuge under 3000 rpm for 10 min, then the supernatant HCl was discarded. The product was washed with ethyl ether twice, then dried under reduced pressure. The product was stored at -30°C until use. (1.44, 91% yield). $^1$H NMR (500 MHz, $D_2O$) δ 3.82 (s, 2H), 3.50 (s, 4H)

*Synthesis of compound **3**: azido-aldehyde*

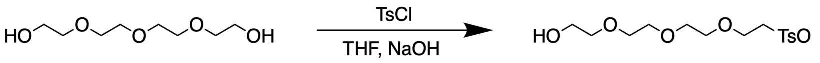

Tetraethylene glycol (50g, 257.44 mmole, 12 eq.) was added to 10 mL of tetrahydrofuran (THF) and a solution of NaOH in 8 mL water (1.37g, 34.32 mmole, 1.6 eq.) was slowly added to a mixture. The mixture was cooled down to 0C in ice bath. Tosyl chloride (TsCl, 4.09g, 21.45 mmole, 1 eq.) in 25 mL THF was added dropwise with stirring using syringe pump (50 mL syringe, diameter = 29.1 mm, rate = 5mL/hr). The system was connected to gas bubbler to release pressure. After stirring at 0°C for 5-6h the mixture was left stirring at RT overnight. The product was extracted with dichloromethane (DCM), washed with water then brine, and dried with sodium sulfate. The solvent was removed under reduced pressure, and **intermediate product 1 (Ts(PEG)$_4$)** remained as sticky solution. (6.16g, 82% yield)

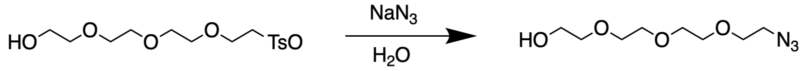

The mixture of **intermediate product 1 (Ts(PEG)$_4$)** (6.157g, 17.69 mmole, 1 eq.) and sodium azide (2.3g, 35.39 mmole, 2 eq.) 41 mL of DI water was stirred at 110°C in oil bath overnight and protected from light. The **intermediate product 2 ((PEG)$_4$-N$_3$)** was extracted with DCM, dried with sodium sulfate, then purified by silica gel chromatography (ethyl acetate). (1.31g, 34% yield)

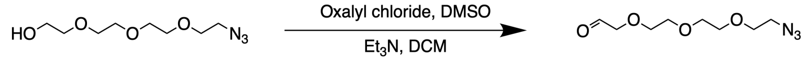

Oxalyl chloride (1.018 mL, 11.87 mmole, 2 eq.) was dissolved in 40 mL of DCM (dry) then cooled down to -78°C in acetone/dry-ice bath. Dimethyl sulfoxide (1.686 mL, 23.74 mmole, 4 eq.) was added dropwise over 10 min then the mixture was stirred for 30 min. The **intermediate product 2 ((PEG)$_4$-N$_3$)** (1.3g, 5.94 mmole, 1 eq.) was dissolved in 3 mL DCM then added dropwise into mixture then stirred for 1 hr. Triethylamine (4.955 mL, 35.62 mmole, 6 eq.) was added dropwise into mixture over 10 min then stirred for 1 hr. The mixture was diluted with DCM, washed with ammonium chloride, water, then brine, and dried with sodium sulfate then the solvent was removed by rotavap. The final product **compound 3 (azido-azide)** was purified by silica gel chromatography (ethyl acetate). (0.588 g, 46% yield). $^1$H NMR (300 MHz, CDCl$_3$) δ 9.74 (t, $J$ = 0.9 Hz, 1H), 4.17 (d, $J$ = 0.9 Hz, 2H), 3.77 – 3.64 (m, 10H), 3.39 (t, $J$ = 5.0 Hz, 2H).

**Supplementary figures.**

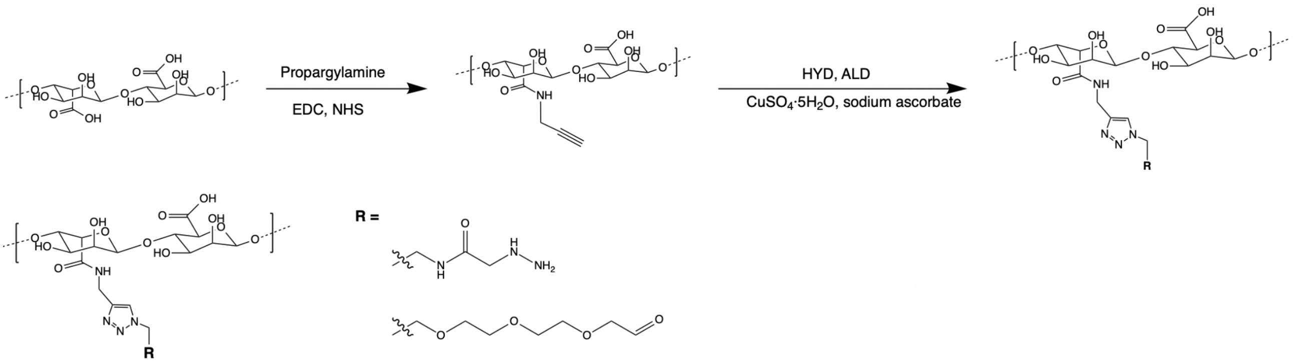


**Figure S1.** Alginate modification scheme. Alginate was first functionalized with alkyne, followed by click reaction with N-(3-azidoethyl)-2-hydrazineylacetamide or azido-aldehyde to obtain alginate-hydrazine (AG-HYD) or alginate-aldehyde (AG-ALD) respectively.

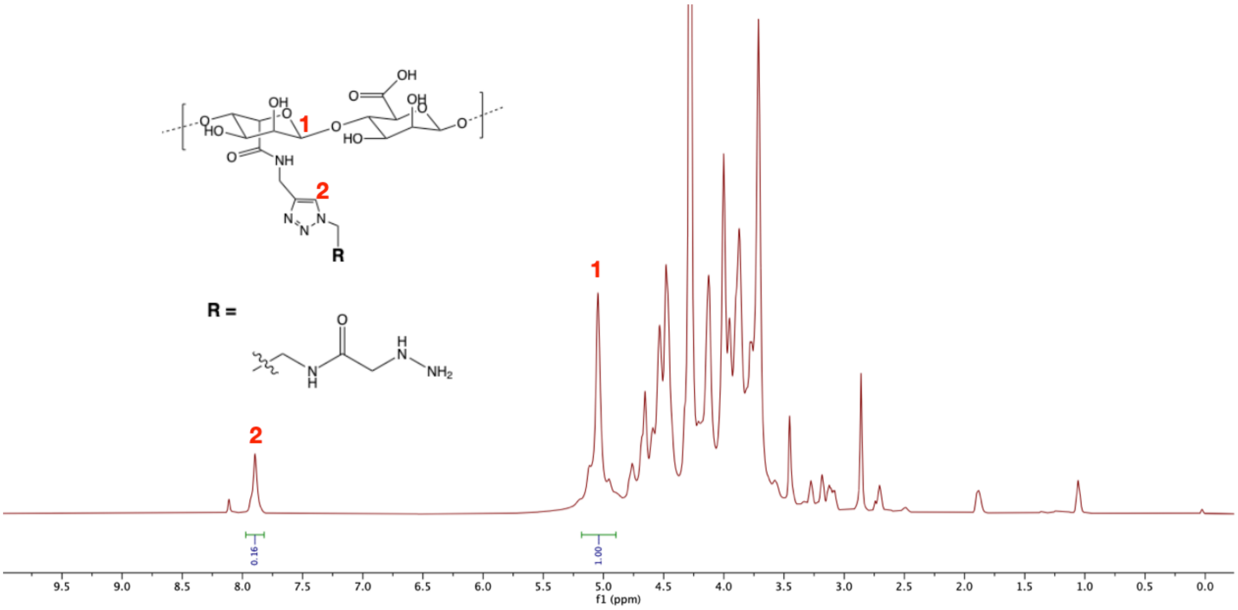


**Figure S2.** $^1$H NMR spectrum (in $D_2O$) of AG-HYD was obtained at 70°C to expose the peak of hydrogen (G1H) on first carbon of guluronic acid (G). The degree of substitution (DS%) of 9.6% was calculated by the ratio of triazole hydrogel (2) to G1H (1) peaks, accounted for the G/M ratio of 1.5, or 60% alginate repeating units are G block.

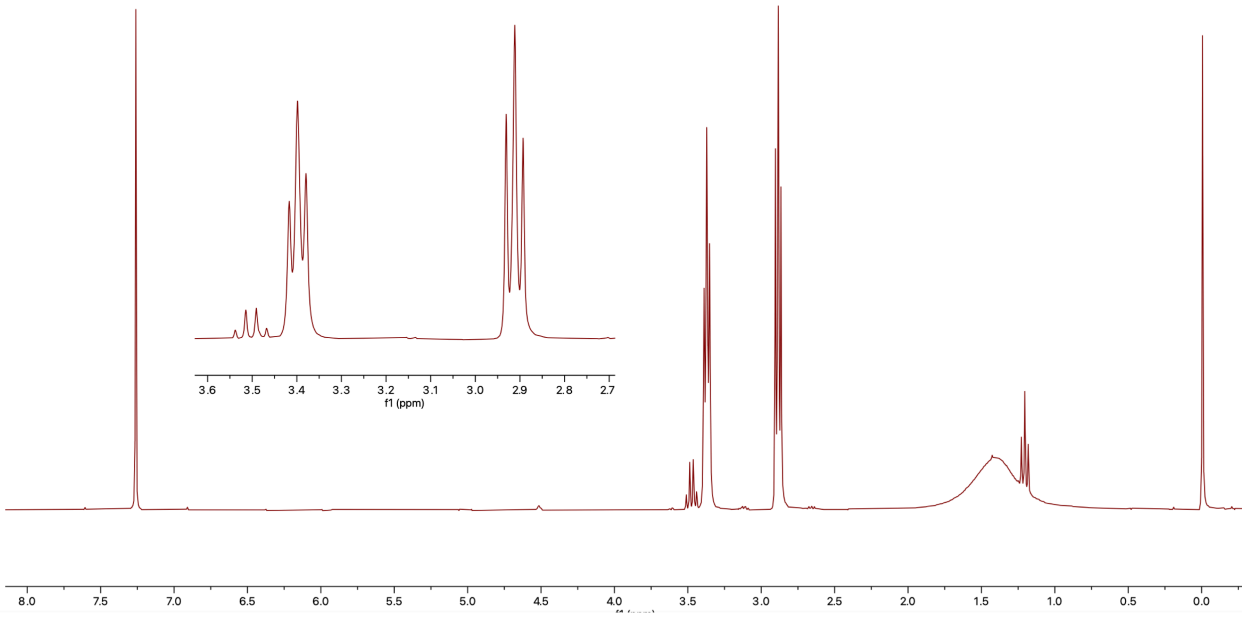


**Figure S3.** $^{1}$H NMR spectrum (in $CDCl_3$) of compound **1**: 2-azidoethylamine

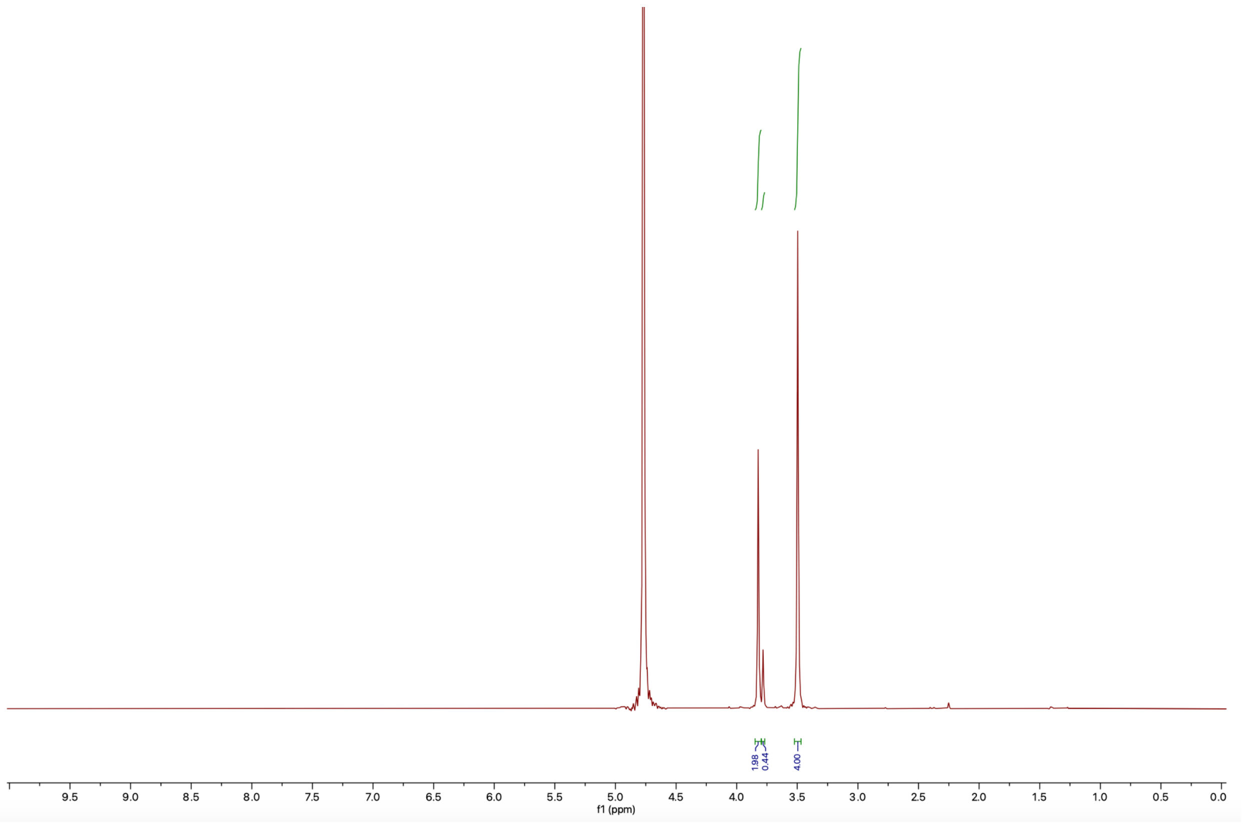


**Figure S4.** $^{1}$H NMR spectrum (in $D_2O$) of compound **2**: N-(3-azidoethyl)-2-hydrazineylacetamide

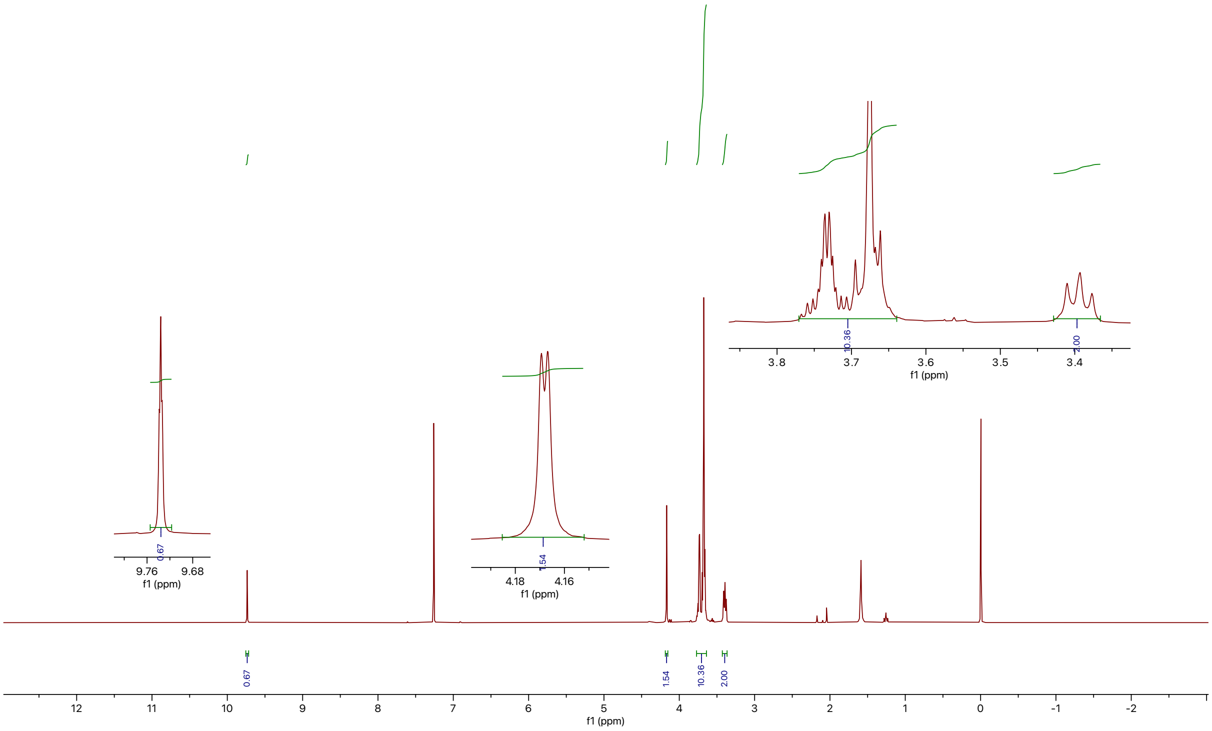


**Figure S5.** $^1$H NMR spectrum (in $CDCl_3$) of compound **3**: azido-aldehyde

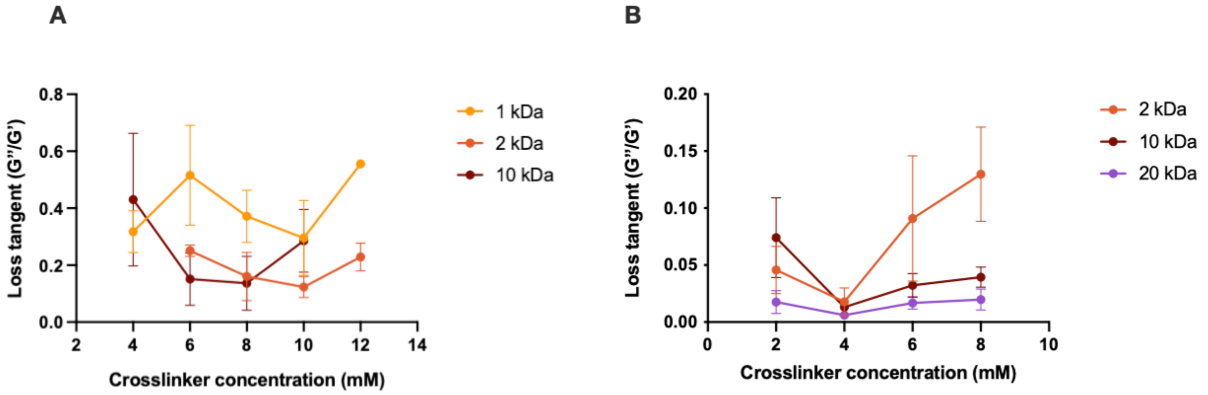


**Figure S6.** The loss tangent at various crosslinker concentration for LX hydrogels (A) and SX hydrogels (B). All samples have AG-HYD of 2 wt%.

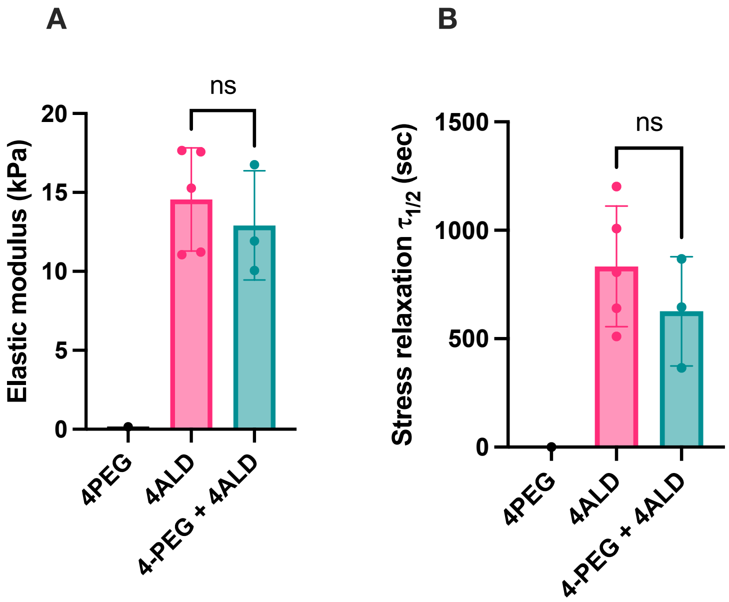


**Figure S7.** High PEG-aldehyde crosslinker concentration does not lead to entanglement in hydrogels. AG-HYD mixed with star crosslinker without aldehyde group (4PEG, 20 kDa) did not form a gel (A). AG-HYD mixed with 4PEG in addition to star crosslinker with aldehyde (4ALD, 20 kDa) did not alter the elastic modulus (A) and stress relaxation time (B) compared to hydrogels with only AG-HYD and 4ALD. Hydrogel formulation: 2wt% AG-HYD (10.2%DS) with 3 mM of 4PEG, 3 mM of 4ALD, or 3 mM of 4PEG + 3 mM of 4ALD. Polymers were dissolved in 0.1M PBS, PH=7.50. One-way ANOVA with Tukey's post-hoc test. ns, not significant.

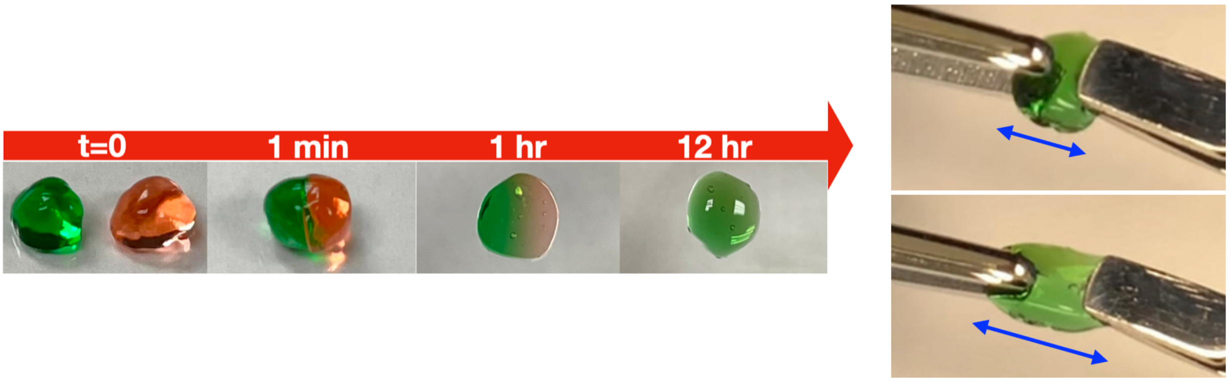


**Figure S8.** Self-healing of SX hydrogels. Two of the star crosslinking hydrogels were dyed with different colors, and gently pressed down with a coverslip with weight to ensure good contact between gels. The different colors of the hydrogels merged over time, and the fused hydrogel did not break when pulling with tweezers, suggesting the hydrogel self-healed. Hydrogel formulation: 2wt% AG-HYD + 4 mM 4ALD (10 kDa) in 0.1 M PBS, pH = 7.50

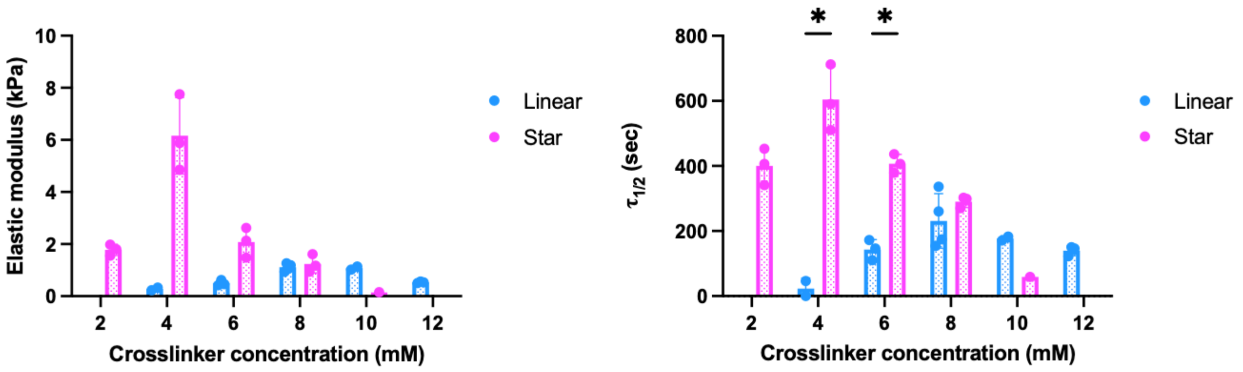


**Figure S9.** SX hydrogels exhibit higher elastic modulus and stress relaxation time compared to LX hydrogels at same crosslinker MW (2kDa). The AG-HYD concentration was kept at 2wt% for all formulations. Multiple unpaired Welch's t-test with Holm-Šídák multiple comparison. ns, not significant; * $P \leq 0.05$.

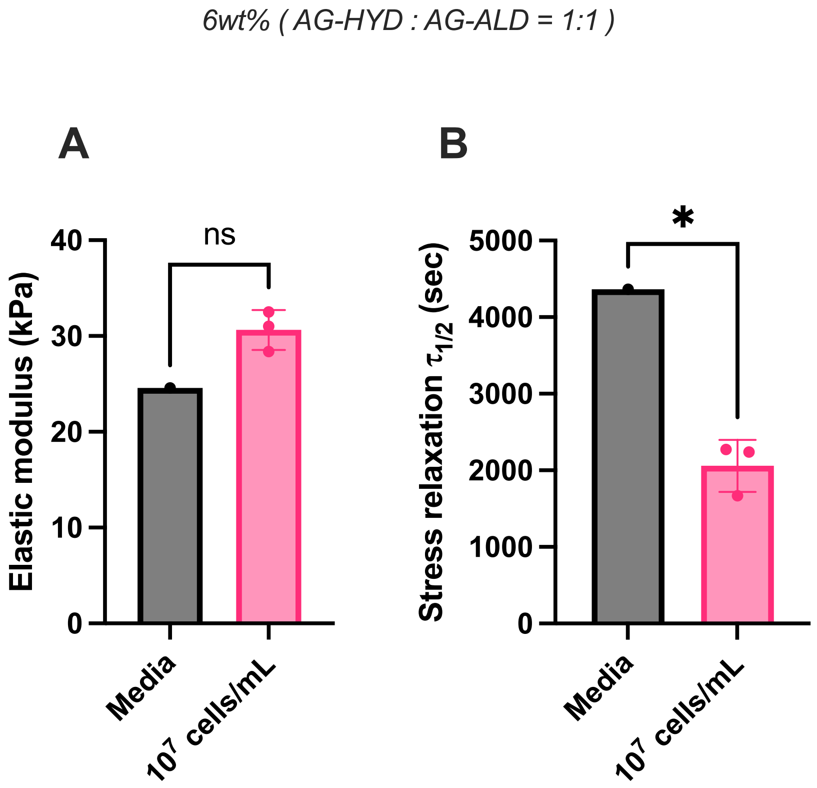


**Figure S10.** The elastic modulus (A) and stress relaxation half-time (B) of SCX hydrogels of 6 wt% alginate with 1:1 AG-HYD to AG-ALD ratio. The mechanical properties were compared with or without encapsulation of 10 million cells per mL in cell culture media. Two-tailed unpaired t-test. ns, not significant; * $P \leq 0.05$.

**Supplementary Table.**

| Polymer | Manufacturer | Catalog number | Batch number | MW (Da) | DS% | Data |
|---|---|---|---|---|---|---|
| Alginate, PRONOVA UP VLVG (AG) | NovaMatrix, IFF | 4200501 | BP-1212-24 | ~28,000 | - | - |
| Alginate-alkyne (AG-ALK) | - | - | AG-ALK-092921 | ~28,000 | 9.6 | - |
| Alginate-hydrazine (AG-HYD) | - | - | AG-HYD-020322 | ~28,000 | 9.6 | All, except Fig. S7 |
| Alginate-hydrazine (AG-HYD) | - | - | AG-HYD-042721 | ~28,000 | 10.2 | Fig. S7 |
| Alginate-aldehyde (AG-ALD) | - | - | AG-ALD-101322 | ~28,000 | 9.6 | Fig. 2, 4, 5, S10 |
| PEG-dialdehyde, 1k Da (2ALD) | Creative PEGWorks | PSB-317 | FY06037 | 1036 | 95 | Fig. 3, 4, S6 |
| PEG-dialdehyde, 2k Da (2ALD) | Creative PEGWorks | PSB-318 | TZQ10114 | 1980 | 95 | Fig. 3, 4, 5, 6, S6, S9 |
| PEG-dialdehyde, 10 kDa (2ALD) | Creative PEGWorks | PSB-322 | GC07082 | 10520 | 95 | Fig. 3, 4, 5, S6 |
| 4-arm PEG-aldehyde, 2 kDa (4ALD) | Creative PEGWorks | PSB-4301 | FY07035 | 2140 | 99 | Fig. 3, 4, 5, S6, S9 |
| 4-arm PEG-aldehyde, 10 kDa (4ALD) | Creative PEGWorks | PSB-4303 | FY08083 | 10140 | 99 | Fig. 3, 4, 5, S6, S8 |
| 4-arm PEG-aldehyde, 10 kDa (4ALD) | SINOPEG | 6020701409 | YP2103070140901 | 9869 | 100 | Fig. 3, 4, 5, S6 |
| 4-arm PEG-aldehyde, 20 kDa (4ALD) | Creative PEGWorks | PSB-4304 | LZ19033 | 21242 | 95 | Fig. 3, 4, 5, S6, S7 |

**Table S1.** Information of polymer used in this study.
